# Autonomous neuronal resilience under metabolic stress highlights fundamental differences between hESC and hiPSC-derived neurons

**DOI:** 10.1101/2025.08.15.670492

**Authors:** SeungJu Yoo, Eunjin Yang, Minee L. Choi

## Abstract

Human induced pluripotent stem cells (hiPSCs) aim to replicate the developmental and functional capacity of human embryonic stem cells (hESCs). Here, we identify neuronal resilience under metabolic stress as a critical parameter for benchmarking equivalence. Without medium replenishment, hESC-derived cortical neurons underwent *neural resurrection*—a spontaneous recovery marked by increased cell density, preserved morphology, and sustained calcium signaling and mitochondrial function for 23 days. In contrast, hiPSC-derived neurons survived but showed reduced adaptability, deteriorating within 10–14 days despite higher initial densities. Our findings introduce a physiologically relevant assay for stress resilience and highlight the need to optimize hiPSC differentiation to achieve hESC-like performance, improving their translational value for disease modeling and regenerative therapy.

## INTRODUCTION

Human induced pluripotent stem cells (hiPSCs) have transformed neuroscience by enabling the derivation of patient-specific neurons for disease modeling, drug discovery, and regenerative therapy (Takahashi et al., 2007; Park et al., 2008). Their accessibility and ethical advantages have positioned hiPSCs as a leading platform for modeling human brain disorders in vitro. However, the ultimate goal for hiPSC-derived neurons is to fully replicate the developmental, functional, and adaptive properties of human embryonic stem cell (hESC)-derived neurons, which remain the gold standard for pluripotency and differentiation (Wernig et al., 2007; Chin et al., 2009; Araki et al., 2013).

While considerable progress has been made in matching the electrophysiological and molecular profiles of hESC-derived neurons, less attention has been given to functional resilience—the capacity of neurons to maintain viability and recover structure and function under prolonged stress. This property is critical in both physiological contexts (e.g., development, plasticity) and pathological states (e.g., injury, neurodegeneration). A systematic comparison of resilience between hESC- and hiPSC-derived neurons is lacking, leaving a gap in our understanding of how closely reprogrammed cells approximate their embryonic counterparts.

Here, we short report neuronal resilience under metabolic stress as a simple but powerful benchmark for stem cell–derived neurons. By subjecting hESC- and hiPSC-derived cortical neurons to nutrient deprivation without medium replenishment, we observed a striking difference in adaptability. hESC-derived neurons exhibited *autonomous neural resurrection*— a spontaneous structural and functional recovery sustained for over three weeks—whereas hiPSC-derived neurons also survived but showed significantly reduced regenerative capacity, with cultures failing within two weeks.

These findings reveal an underappreciated functional gap between hESC- and hiPSC-derived neurons, highlight cell-origin–dependent adaptability as a critical performance metric, and provide a physiologically relevant assay for optimizing hiPSC differentiation protocols to achieve hESC-like robustness for disease modeling and therapeutic use.

## RESULTS

### Observation under cellular fasting in hESC-derived neurons

hESCs were differentiated into cortical neurons and verified by morphological assessment (data not shown; see Fig. 1A for the differentiation timeline). Cells were seeded at low density, resulting in limited cell–cell contact—conditions typically unfavorable for neuronal survival. At Day 0, shortly after passaging onto new chambered slides, neurons exhibited fragmented processes, irregular somata, and poor network connectivity.

**Figure 1.**
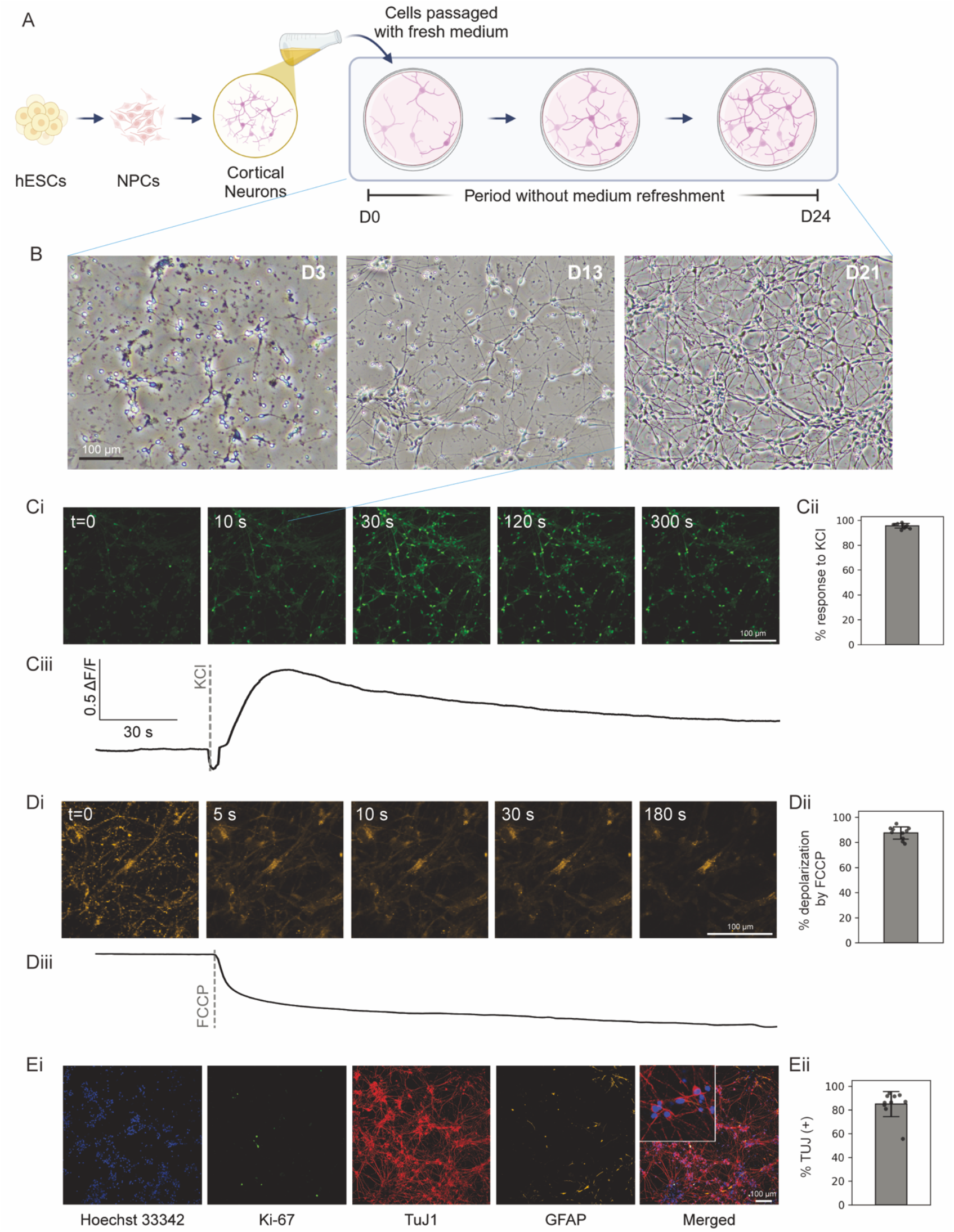
hESC-Derived Neurons Exhibit Resilience and Functional Maturation During 23 Days of Cellular Fasting. (A) Schematic timeline illustrating the observation of hESC-derived neurons under prolonged metabolic stress without medium replenishment (*cellular fasting*). (B) Phase-contrast images showing morphological progression from initial unhealthy appearance at Day 0 to substantial recovery and network formation at Day 3, Day 13, and Day 21. (C) Functional validation by calcium imaging at Day 18. (Ci) Representative images and (Cii) traces showing robust responses to KCl stimulation, indicating the presence of potassium-dependent ion channels. Calcium imaging was performed using the Fluo-4 intracellular calcium indicator. (D) Mitochondrial membrane potential assessment using TMRM, with FCCP applied as a mitochondrial uncoupler. (E) Immunocytochemistry confirming that the majority of cells were TuJ1^+^ neurons (>85%), with small fractions of Ki-67^+^ proliferating cells (<1%) and GFAP^+^ glial cells (<5%). GFAP^+^ cells were primarily located in lower culture layers.

Over the following 24 days without medium replenishment (*cellular fasting*), cultures underwent a striking and progressive recovery (Fig. 1B). Neuronal morphology became more regular, dendritic and axonal processes extended extensively, and cell density increased, culminating in a dense, interconnected neural network. The observation ended at Day 24 due to medium evaporation and exposure to air.

Functional assessment at Day 18 confirmed that these recovered neurons remained active. Calcium imaging revealed that >90% of cells exhibited ≥20% increases in ΔF/F in response to KCl stimulation (Fig. 1C, Cii), indicating preserved excitability. Mitochondrial membrane potential, assessed using TMRM, was also maintained (Fig. 1D). Minimal cell death was observed even 18 days after the last medium change, indicating a high degree of resilience and sustained viability under prolonged metabolic stress.

Immunocytochemistry demonstrated that the cultures were composed predominantly of TuJ1^+^ neurons (>85%), with minor populations of GFAP^+^ glial cells (<5%) and Ki-67^+^ proliferating cells (<1%) (Fig. 1E). GFAP^+^ cells were primarily localized to the lower culture layers. ATP stimulation elicited no detectable calcium response (Supplementary Fig. 1A), consistent with the absence of mature astrocytes.

### hiPSC-Derived Neurons Exhibit Limited Resilience Compared to hESC-Derived Neurons

To assess whether the *neural resurrection* phenomenon observed in hESC-derived neurons also occurred in hiPSC-derived neurons, we performed parallel experiments under identical nutrient-deprivation (*cellular fasting*) conditions. At the onset of medium withdrawal, hiPSC-derived neurons maintained initial morphology; however, signs of cellular stress appeared within days, including medium discoloration consistent with pH shifts, increased cell death, disrupted neuronal architecture, and the proliferation of non-neuronal cell types.

Even when plated at higher initial densities to promote cell–cell contact and network formation, hiPSC-derived neurons underwent progressive degeneration, with cultures failing to survive beyond 10–14 days (Fig. 2A, B). Unlike their hESC-derived counterparts, these cultures did not display morphological recovery or network re-establishment over time.

**Figure 2.**
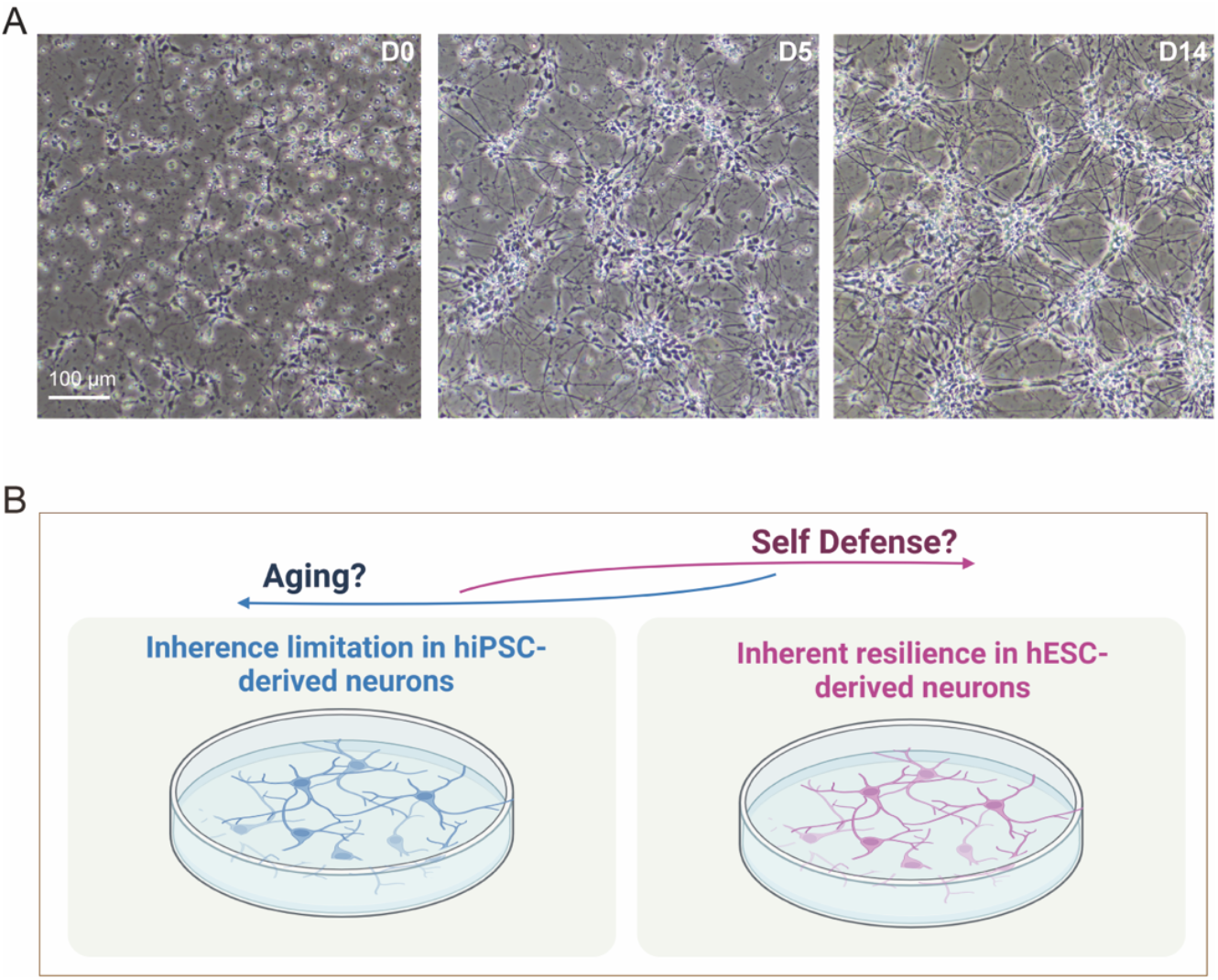
hiPSC-derived neurons remain morphologically healthy for up to 14 days without medium replenishment (A), with the overall experimental design illustrated schematically in (B).

Repeated inability to reproduce the *neural resurrection* effect in hiPSC-derived neurons was consistently associated with medium acidification, whereas hESC-derived cultures maintained pH stability. This suggests that extracellular pH homeostasis may be a critical determinant of prolonged neuronal survival under metabolic stress. Reduced resilience in hiPSC-derived neurons therefore reflects a diminished capacity to regulate their extracellular environment, identifying pH regulation as both a functional readout of neuronal robustness and a potential target for optimizing hiPSC differentiation protocols.

## DISCUSSION

In vitro models of the human brain offer exceptional potential for advancing our understanding of nervous system complexity and function. These platforms enable the study of intricate cellular interactions, synaptic dynamics, and disease mechanisms in a controlled setting, providing unparalleled insights into both normal physiology and pathological processes (Connors et al., 1982; Okabe et al., 1996; van Pelt et al., 2005). Among these, human induced pluripotent stem cells (hiPSCs) have emerged as a transformative tool due to their derivation from patient-specific somatic cells. This capability allows non-invasive investigation of the genetic and molecular basis of individual disorders, opening opportunities for personalized medicine and disease modeling (Jafari, 2023; Kim et al., 2021; Virdi et al., 2022).

However, the conversion of hiPSCs into fully functional neurons remains challenging, particularly when compared with neurons derived from human embryonic stem cells (hESCs). While hiPSCs are valued for their accessibility and ethical advantages, intrinsic differences between hESCs and hiPSCs may influence differentiation potential and resilience. These differences often become more pronounced under extreme or suboptimal culture conditions, where hESC-derived neurons frequently display greater adaptability, resilience, and neurogenic capacity than their hiPSC-derived counterparts.

In this report, hESC-derived neurons demonstrated remarkable resilience under *cellular fasting*—surviving for 23 days without medium replenishment while maintaining the ability to generate structurally intact, functional neurons. In contrast, hiPSC-derived neurons exhibited significant stress responses, including medium acidification, increased cell death, and loss of normal morphology, with survival limited to 10–14 days even at higher initial seeding densities. This highlights both the superior adaptability of hESC-derived neurons and the resilience limitations of current hiPSC-derived neuronal models.

Such intrinsic differences in robustness have direct implications for the refinement of neural differentiation protocols. Understanding and addressing the limitations of hiPSC-derived neurons is essential to improving their reliability in disease modeling, regenerative medicine, and the production of functional neural tissues equivalent to those derived from hESCs.

Sufficient nutrition. It is likely that residual nutrients in the culture medium were sufficient to sustain neuronal survival for the duration of observation, in contrast to standard protocols requiring medium changes every 3–5 days. To examine the role of initial medium composition, we replaced N2B27 with DMEM/F-12 supplemented with GlutaMAX, which lacks neuron-specific supplements. Neurons maintained in DMEM with routine medium renewal for up to 14 days showed strong calcium responses (Supplementary Fig. 1A) and healthy mitochondrial membrane potential. However, their resilience was reduced compared to N2B27-preconditioned neurons under fasting, surviving no longer than two weeks. This supports the hypothesis that preconditioning with N2B27 contributed to prolonged survival during fasting.

Are hESC-derived neurons still “young”? hESC-derived neurons are generally considered “young” in cellular age, a state that likely contributes to their enhanced resistance to stress and injury—features typically observed in immature neurons. Consequently, they may more closely resemble the cellular and functional characteristics of a young brain. In contrast, hiPSC-derived neurons may retain epigenetic or molecular hallmarks of their donor’s chronological age, despite reprogramming. While reprogramming resets many cellular features, it may not completely erase donor age–associated epigenetic memory (Kim et al., 2010; Samoylova & Baklaushev, 2020). As a result, hiPSC-derived neurons may exhibit properties more characteristic of aged neurons. This differential resilience could be exploited for modeling age-related neurodegenerative diseases, such as Parkinson’s disease, where vulnerability to stress is a defining feature.

Altered internal state for self-defense. Under prolonged nutrient deprivation, neurons may activate adaptive survival programs. Detection of stress signals from compromised neurons can trigger gene expression changes that promote cell health, secretion of neurotrophic factors, and modulation of energy metabolism to reduce stress burden (Bar-Ziv et al., 2023; Fulda et al., 2010; Kourtis & Tavernarakis, 2011; Munns et al., 2003). These adaptations may also involve remodeling of the extracellular matrix and modification of the local microenvironment to support culture-wide survival.

Conclusion. Even without medium replenishment, hESC-derived neurons demonstrated extraordinary adaptability, maintaining viability, morphology, and function over extended periods. This adaptability likely fosters a favorable microenvironment that supports ongoing neuronal survival and network integrity. Such resilience reflects the intrinsic regenerative capacity of hESC-derived neurons and underscores resilience as a defining performance metric. For hiPSC-derived neurons to fully meet the functional standards set by hESC-derived counterparts, achieving comparable resilience will be essential. This distinction highlights the importance of investigating the mechanisms governing neuronal adaptability, with direct relevance to disease modeling and regenerative applications.

**Supplementary Figure 1.**
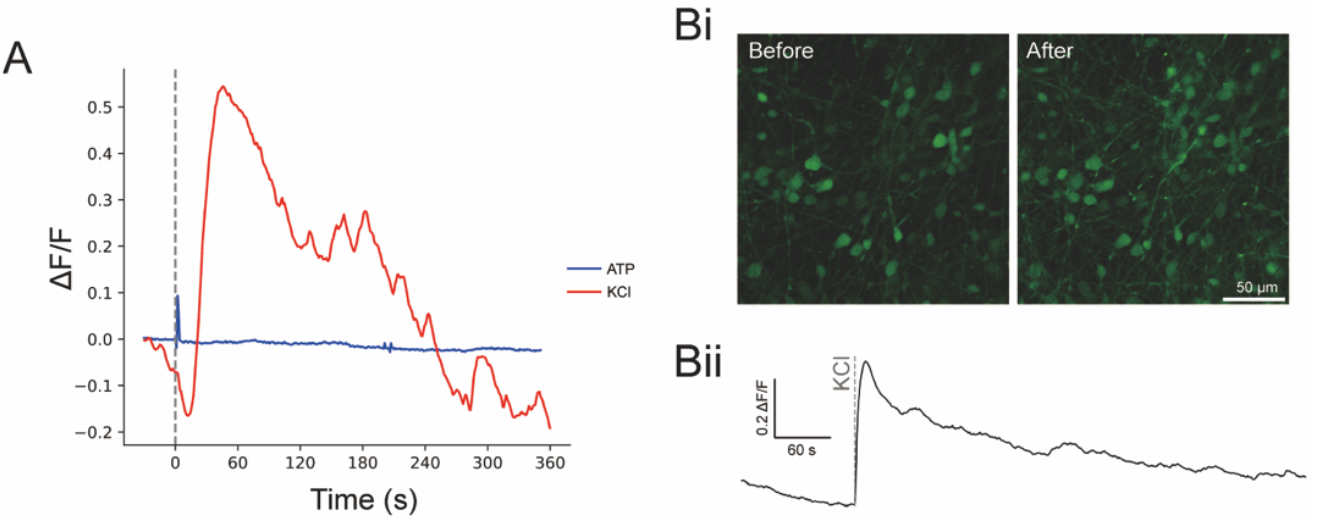
Neurons do not respond to ATP stimulation. Representative calcium imaging traces (A) and corresponding images (B) are shown.

## METHODS

### Stem cell maintenance

All cell lines were cultured based on the protocol provided by the supplier. The H9 human embryonic stem cell (hESC) line was gifted by Ki-Jun Yoon’s laboratory (Department of Biological Sciences at KAIST). The SCTi-003A human induced pluripotent stem cell (hiPSC) line was obtained from STEMCELL Technologies. Specific information about stem cell lines is included in Table 1. H9 was cultured on Matrigel-coated plates with StemFlex medium. SCTi-003A was cultured with mTeSR plus medium. They were considered to have reached 60% confluency when stem cell colonies began to merge. When the confluency exceeded 60%, the cells were passaged using 0.5mM EDTA diluted in DPBS. All feeder-free stem cells were maintained in a 5% CO_2_ incubator at 37°C and confirmed to be mycoplasma-free by Cosmogenetech. Details of all materials and reagents used are listed in Table 2.

**Table 1.**
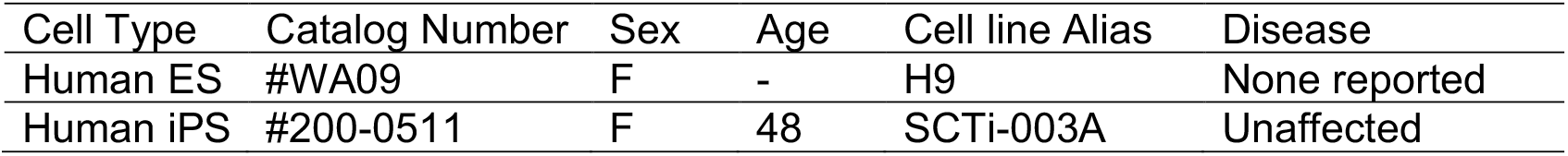
Characteristics of Stem Cell Line.

**Table 2.**
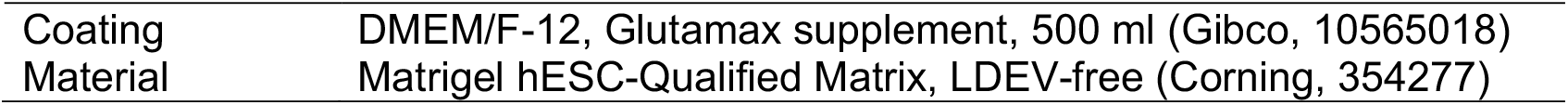
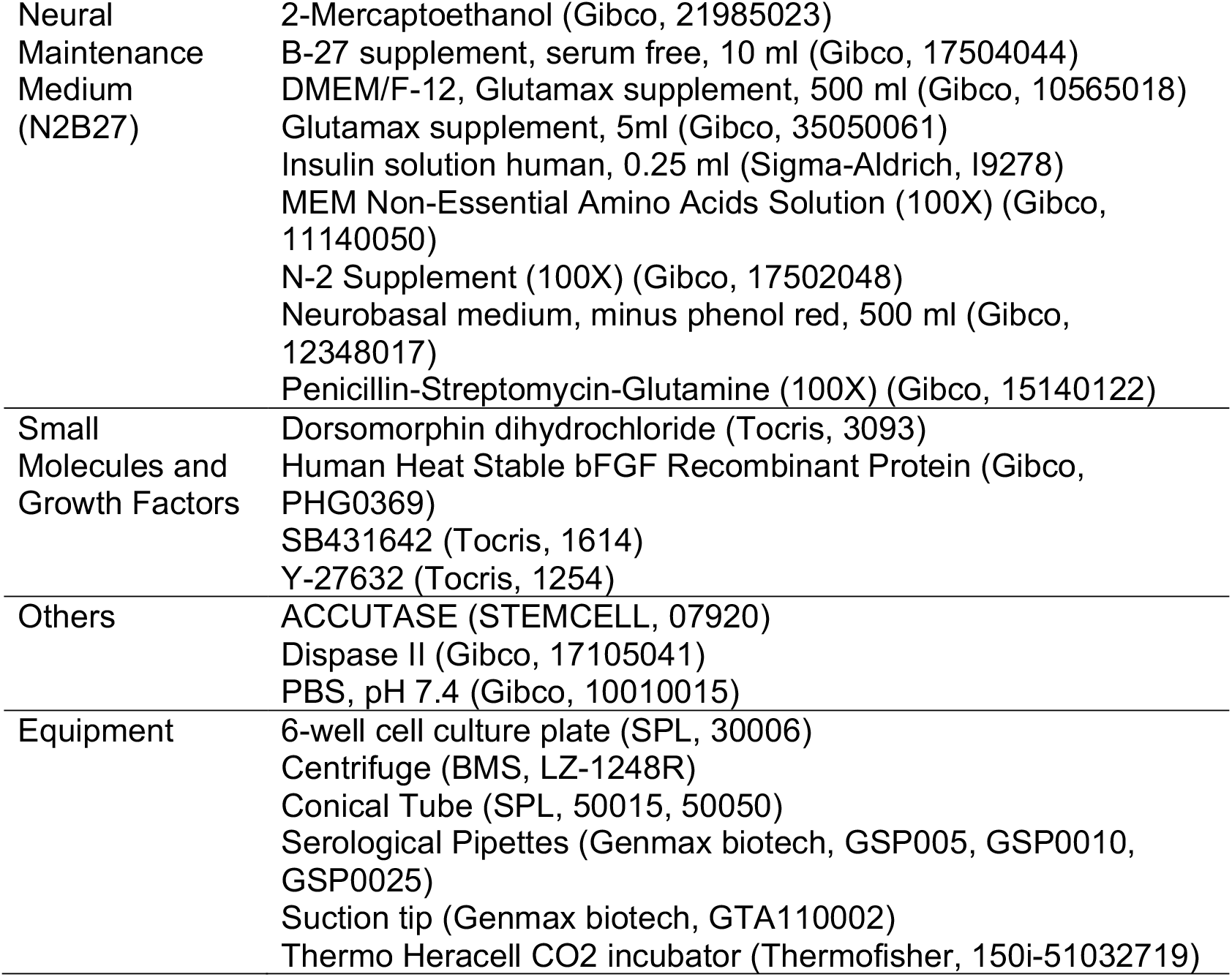
Materials for Stem cell maintenance and Cortical neuron differentiation.

### Cortical neuron differentiation

For cortical neuron differentiation, the protocol by Shi et al. (2012) was modified. When the cells were grown to over 95% confluency, the stem cell maintenance medium was replaced with N2B27 medium (a 1:1 mixture of DMEM/F12 with Glutamax and Neurobasal without phenol red, 0.5% (v/v) N2 supplement, 1% (v/v) B27 supplement, 0.5% (v/v) MEM-NEAA, 0.5% (v/v) Glutamax, 0.025% (v/v) Insulin, 0.1% β-mertoethanol, and 0.5% Penicillin/Streptomycin) supplemented with 10 µM SB 431642 and 1 µM Dorsomorphin dihydrochloride. Neural induction media was replaced daily for 10 days to induce neuroepithelium via dual SMAD inhibition. When cells formed a single epithelial sheet around day 10, they were detached using 1.8U/mL Dispase and plated on a Matrigel-coated plate with N2B27 medium supplemented with 10 µM Y-27632 at a 1:2 ratio. The medium was replaced every other day. Although there were slight differences between cell lines, neural rosettes typically began to appear around day 20. After day 20, cells were maintained at appropriate density with passages using Accutase. As noted by Shi et al. (2012), the cortical neural progenitor cell population exceeded 99% around day 28 and late cortical neuron population developed around day 80. Details of all materials and reagents used are listed in Table 2.

### Immunocytochemistry

To prepare the sample for immunocytochemistry, µ-Slide 8-well glass bottom chambered slide was coated with Matrigel. Differentiated neurons were detached using Accutase, counted, and plated at a density of 150,000 cells per well, ensuring the total media volume per well did not exceed 250 µl. Y-27632 was supplemented in the media depending on the condition of the cells.

When the cells were ready for imaging, they were washed once with PBS, and incubated with 170 µl of 4% paraformaldehyde (PFA) per well at room temperature for 40 minutes. After fixation, the wells were washed three times with PBS to remove residual PFA. To validate the identity of cells, the fixed samples underwent permeabilization, blocking, and loading of primary and secondary antibodies. For permeabilization, the fixed cells were loaded with 200 µL of 0.2% Triton X-100 diluted in PBS and incubated at room temperature for 3-5 minutes. Following permeabilization, 0.2% Triton X-100 was removed, and blocking was performed by adding 5% BSA diluted in PBS for 1 hour at room temperature. After blocking, the primary antibody was directly loaded without washing. The primary antibody was prepared at a 1:250 dilution in a 1% BSA solution and applied for either 1 hour at room temperature or overnight at 4°C. After primary antibody incubation, the wells were washed three times with 1% BSA (5 minutes each) for 5 minutes each to reduce non-specific binding. Secondary antibody solution was then loaded at a 1:500 dilution in 1% BSA. As the secondary antibody was fluorescently labeled, the dye dilution and loading were performed in the dark. Secondary antibody incubation was conducted at room temperature for 1 hour. Following tagging, the cells were washed once with 1% BSA for 5 minutes, then washed with Hoechst solution for 15 minutes. Hoechst solution was prepared as 10 µM Hoechst in either PBS or 1% BSA. Finally, a 5-minute wash with PBS was performed. The samples were wrapped in foil to protect from light and stored at 4°C. Imaging was conducted using a Nikon AX-R microscope. Details of all materials and reagents used are listed in Table 3.

**Table 3.**
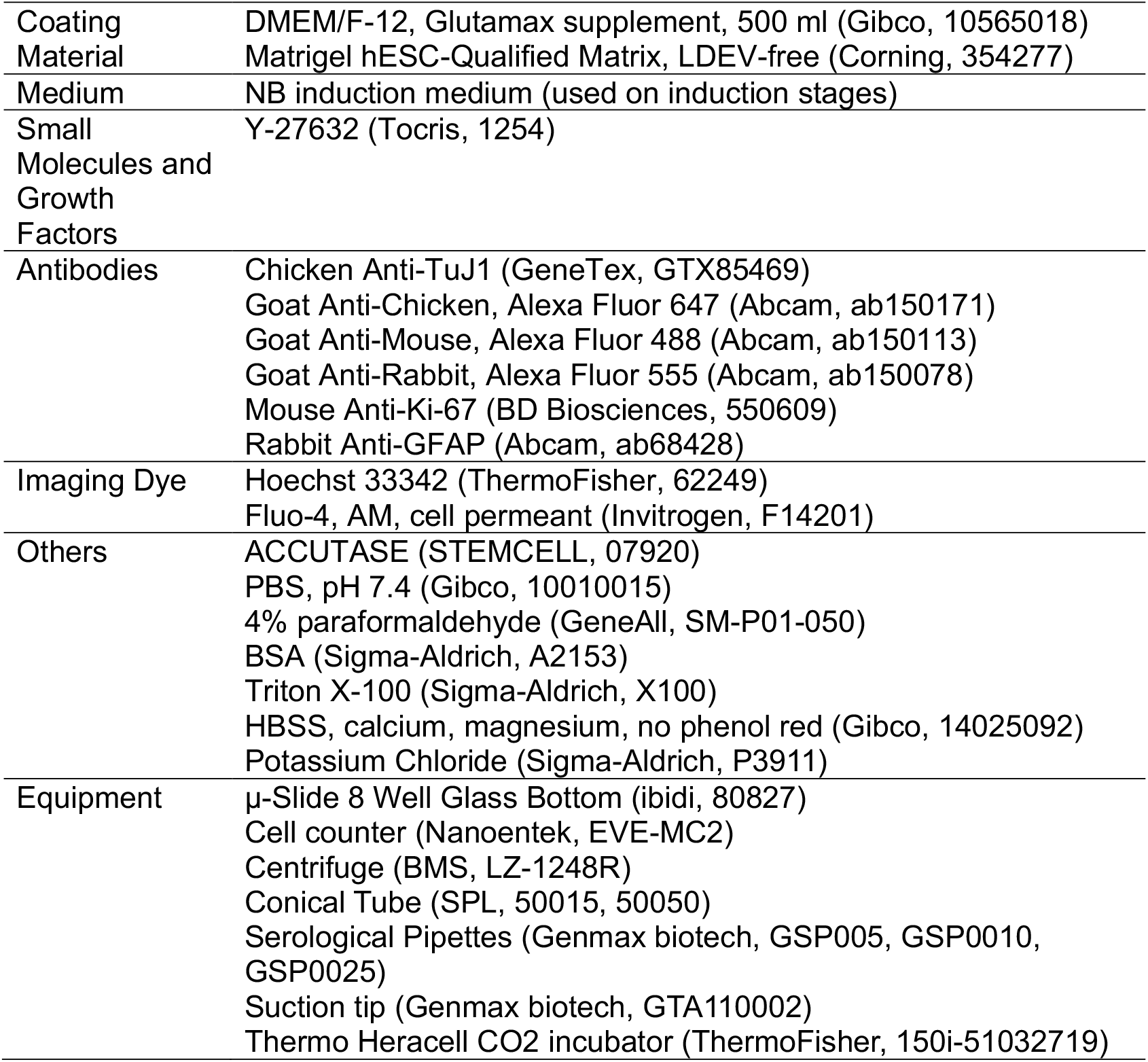
Materials for Immunochemistry, Live Cell Imaging and Calcium Imaging.

### Live Cell Imaging

For live imaging of cortical neurons, sample preparation follows the protocol described in immunocytochemistry part. Live cell imaging was performed to assess organelle function, such as mitochondrial and lysosomal activity, as well as cell viability. Samples plated on the imaging plate were used. Once the samples were ready to be imaged, the medium was replaced with HBSS with calcium and magnesium to maintain cell viability without the need for incubator conditions (5% CO_2_, 37°C). A multiplex dye solution was then prepared, consisting of 10 µM Hoechst for cell nucleus, 25 µM TMRM for mitochondria, 100 nM LysoTracker for lysosome, and 100 µM Sytox for dead cell, all diluted in HBSS. The dyes were prepared fresh, protected from light, and stored according to manufacturer recommendations. Cells were washed once with HBSS, followed by the addition of the multiplex dye solution. The plate was incubated at room temperature in the dark for 30 minutes to stabilize fluorescence signals. After the 30-minute stabilization, imaging was performed using a Nikon microscope AX-R. Details of all materials and reagents used are listed in Table 3.

### Calcium Imaging

To validate functionality of cortical neurons, calcium imaging was performed using Fluo-4 dye and potassium chloride (KCl). Fluo-4 dye, a calcium indicator, was used to detect calcium ion fluctuations, while KCl was applied to induce an action potential-like response in neurons. This approach enabled observation of calcium ion influx and efflux, simulating action potential dynamics in the neurons.

Samples plated on the imaging plate were washed once with HBSS. Cells were then loaded with 5 µM Fluo-4 in HBSS and incubated at room temperature for 40 minutes. After staining, the cells were washed three times with PBS. Following the final wash, fresh PBS was added at 200 µL per well, and imaging was performed using a Nikon AX-R confocal microscope. Imaging was conducted as a time series for approximately 5 minutes, with 25 mM potassium chloride diluted in HBSS loaded during imaging to induce calcium response via membrane depolarization. Imaging was performed using a Nikon microscope AX-R at 1 fps. Details of all materials and reagents used are listed in Table 3.

## Supporting information

Supplementary Figure 1. Neurons do not respond to ATP stimulation. Representative calcium imaging traces (A) and corresponding images (B) are shown

## ACKNOWLEDGEMENTS

We thank the donors for providing fibroblast samples, and we are grateful to Prof. Ki-Jun Yoon for generously sharing the H9 cell line. We also thank Seohyun Kim for assistance with H9 cell line maintenance. This work was supported by the National Research Foundation of Korea, funded by the Korean government (RS-2023-00266872, RS-2024-00343012).

