## Supplementary figures and images for "Autonomous neuronal resilience under metabolic stress highlights fundamental differences between hESC and hiPSC-derived neurons"

### Supplementary Figure 1. Neurons do not respond to ATP stimulation. Representative calcium imaging traces (A) and corresponding images (B) are shown

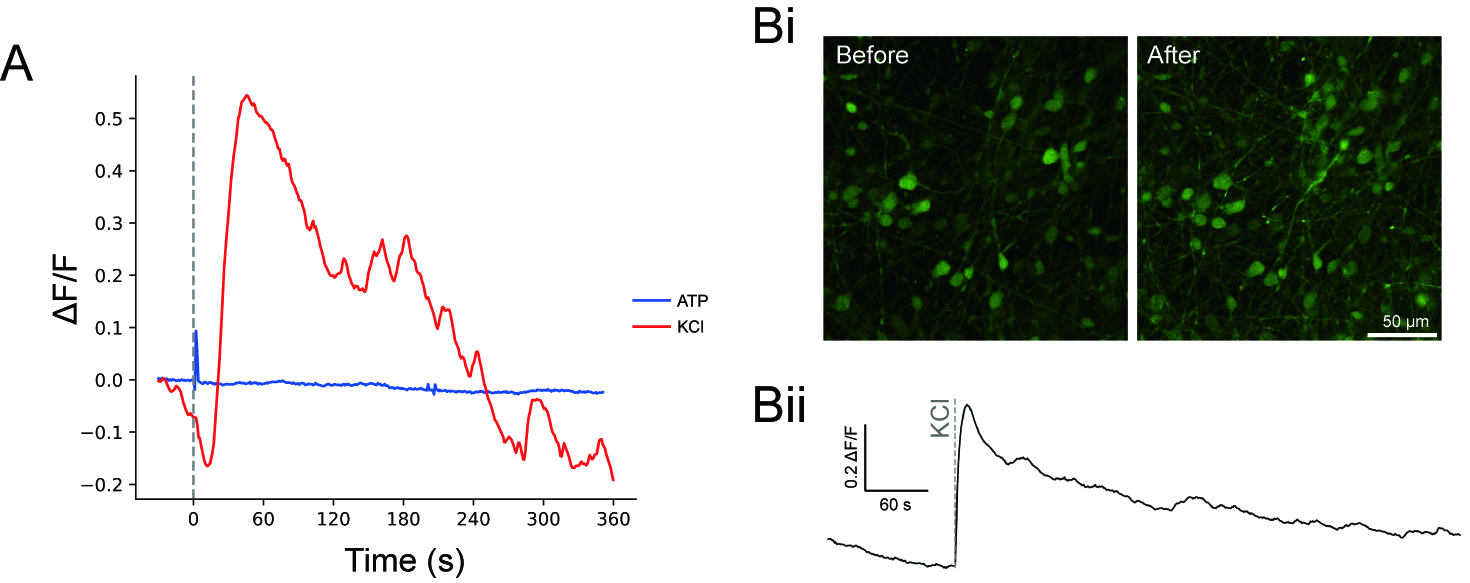
